# EMBL2checklists: A Python package to facilitate the user-friendly submission of plant DNA barcoding sequences to ENA

**DOI:** 10.1101/435644

**Authors:** Michael Gruenstaeudl, Yannick Hartmaring

## Abstract

**Background:** The submission of DNA sequences to public sequence databases is an essential, but insufficiently automated step in the process of generating and disseminating novel DNA sequence data. Despite the centrality of database submissions to biological research, the range of available software tools that facilitate the preparation of sequence data for database submissions is low, especially for sequences generated via plant DNA barcoding. Current submission procedures can be complex and prohibitively time expensive for any but a small number of input sequences. A user-friendly software tool is needed that streamlines the file preparation for database submissions of DNA sequences that are commonly generated in plant DNA barcoding.

**Methods:** A Python package was developed that converts DNA sequences from the common EMBL and GenBank flat file formats to submission-ready, tab-delimited spreadsheets (so-called “checklists”) for a subsequent upload to the public sequence database of the European Nucleotide Archive (ENA). The software tool, titled “EMBL2checklists”, automatically converts DNA sequences, their annotation features, and associated metadata into the idiosyncratic format of marker-specific ENA checklists and, thus, generates output that can be uploaded via the interactive Webin submission system of ENA.

**Results:** EMBL2checklists provides a simple, platform-independent tool that automates the conversion of common plant DNA barcoding sequences into easily editable spreadsheets that require no further processing but their upload to ENA via the interactive Webin submission system. The software is equipped with an intuitive graphical as well as an efficient command-line interface for its operation. The utility of the software is illustrated by its application in the submission of DNA sequences of two recent plant phylogenetic investigations and one fungal metagenomic study.

**Discussion:** EMBL2checklists bridges the gap between common software suites for DNA sequence assembly and annotation and the interactive data submission process of ENA. It represents an easy-to-use solution for plant biologists without bioinformatics expertise to generate submission-ready checklists from common plant DNA sequence data. It allows the post-processing of checklists as well as work-sharing during the submission process and solves a critical bottleneck in the effort to increase participation in public data sharing.

## Introduction

Only a few software tools assist in the preparation of DNA sequence data for submission to public sequence databases, despite the centrality of this process for generating and disseminating novel biological data. Contemporary biological research depends on the preservation, curation, and reproducibility of the data under study [1,2], and the submission of analyzed data to publicly accessible databases constitutes one of the most important best-practices in biology [3,4], particularly in the era of big data [5]. DNA sequences generated to identify and characterize novel organisms or unchartered biodiversity must typically be submitted to public sequence databases before publication of the research is granted [6,7]. Compliance with this prerequisite remains mixed [8–10]. Several large nucleotide sequence repositories accept DNA sequence submissions, including GenBank [11], the European Nucleotide Archive [12] or the DNA Data Bank of Japan [13]. These repositories coordinate their policies and operations under the umbrella of the International Nucleotide Sequence Database Collaboration (INSDC; [14]), but each database employs custom procedures for sequence upload and data submission. ENA, for example, channels the submission of annotated DNA sequences through the Webin submission framework (https://www.ebi.ac.uk/ena/submit/sra/; [15]), which, in its interactive version, operates with pre-formatted, tab-delimited spreadsheets (called “annotation checklists” or “templates”) that are filled out by the user and then uploaded for submission. In order to account for different types of annotated DNA sequences (e.g., coding vs. non-coding, nuclear vs. organellar origin), a series of pre-tailored spreadsheets (hereafter “checklists”) was developed by ENA, each with its idiosyncratic, tab-delimited fields of information. Users of ENA must choose the correct spreadsheet for their data submission (Fig. 1), and different types of DNA sequences must be submitted via separate data uploads. Since June 2017, the submission process through Webin has been automated and now includes automatic validation procedures for annotation features, taxonomic metadata, and sequence integrity [12]. Despite the centrality of data sharing to biological research, the range of user-friendly software tools that assist in data preparation for database submission is perceived as low [3]. Indeed, very few contemporary, user-friendly software tools exist (e.g., Geneious [16]) that facilitate the preparatory steps of annotated DNA sequences prior to uploading them to public sequence repositories. The number of software tools that assist with, and are specifically customized for, the preparation of common plant DNA barcoding sequences is particularly sparse.

**Fig 1.**
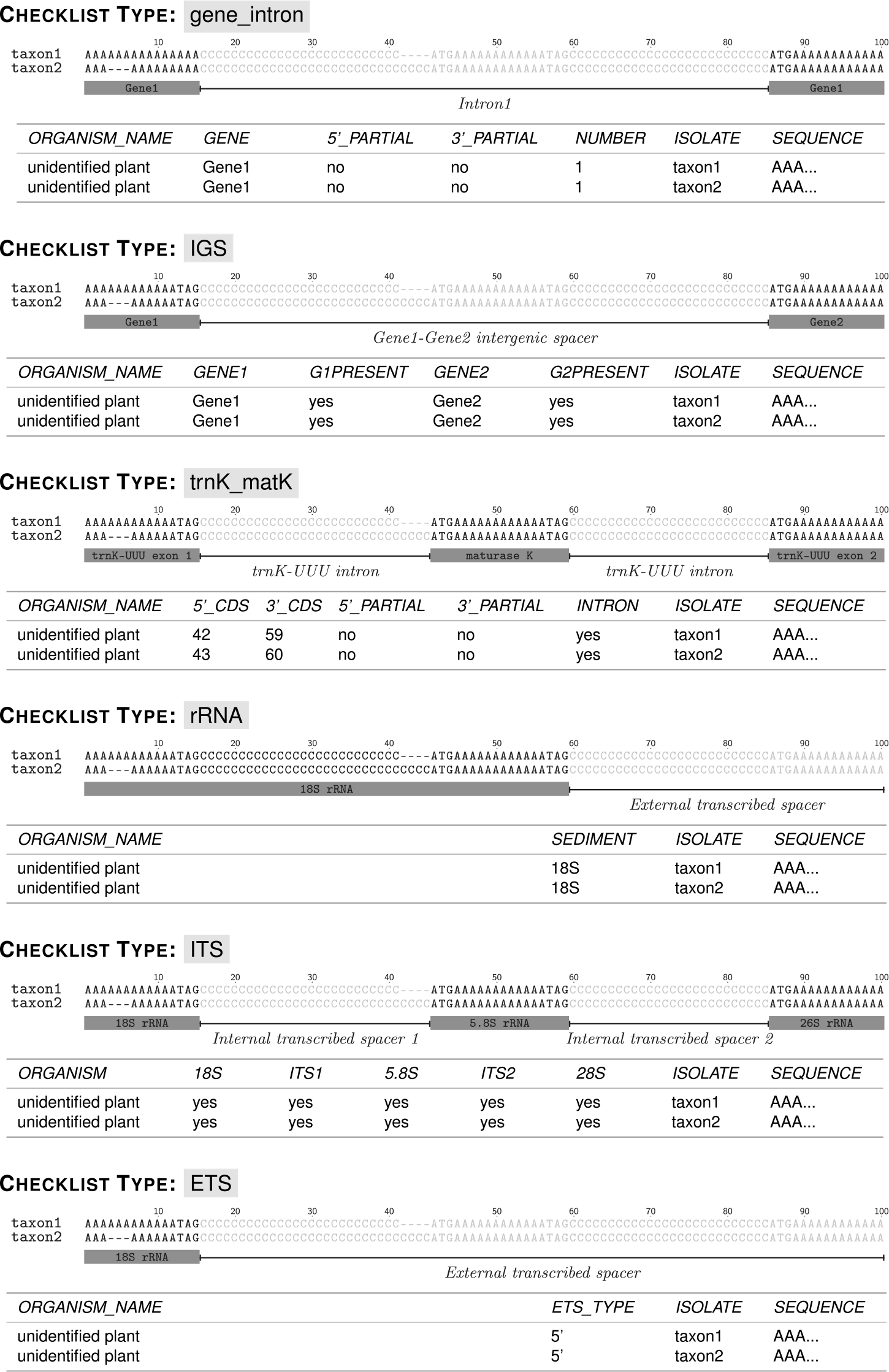
Structural overview of the six DNA markers and their corresponding Webin checklists as implemented in EMBL2checklists. The overview displays the structural characteristics of each DNA marker and the idiosyncratic column names and numbers of the corresponding checklists. The DNA sequences displayed represent dummy sequences and are identical to the sequences of the example test files co-supplied with the software.

Unlike sequence submissions to GenBank, the preparation of DNA sequence data for submission to ENA is insufficiently facilitated, highlighting the demand for software that converts annotated DNA sequences into submission-ready checklists. Upon DNA sequencing, researchers often utilize user-friendly software suites such as Artemis [17], Geneious or PhyDE [18] for the assembly and annotation of DNA sequences. Some of these suits (e.g., Artemis, Geneious) enable the conversion of annotated DNA sequences to file formats that are easily submittable to GenBank, either by producing files in a direct submission format (i.e., the Sequin format [19]) or through the processing with additional tools that convert GenBank-formatted flat files into the Sequin file format (e.g., GB2sequin [20]). While there are similar software suites that generate EMBL-formatted flat files (e.g., Artemis, the “seqret” tool of EMBOSS [21]), they do not produce output files that are suitable for a direct upload and submission via the interactive Webin submission system of ENA. To the best of our knowledge, no conversion tool currently exists that automatically converts annotated DNA sequences and associated metadata from the EMBL flat file format into submission-ready Webin checklists. The Webin submission system is the default gateway for DNA sequence submission to ENA and offers two routes for data upload: an interactive route, in which checklists are uploaded or generated online; and a programmatic route, in which both checklists and pre-formatted flat files can be submitted to the ENA server [12]. Flat files in EMBL file format are not currently accepted through the interactive, but only through the programmatic Webin submission route. However, using the programmatic route requires bioinformatics expertise and is, thus, inaccessible to many users. In absence of a more suitable solution, many users are compelled to undergo the tedious process of parsing information from EMBL-formatted flat files and copying them into Webin checklists manually, or to submit their data to repositories with a richer toolkit for submission preparation (e.g., GenBank). Some users also employ the guided web interface of the Webin submission platform, in which users click their way through an elaborate data entry interface. For any but a small number of input sequences, the use of the guided web interface is prohibitively time-expensive. Thus, there is a strong demand for an easy-to-use, platform-independent software tool that converts DNA sequences in EMBL flat file format including their annotation features and associated metadata into submission-ready checklists for upload via the Webin submission system.

Plant DNA barcoding is a key method to identify and characterize botanical specimens, and the resulting DNA sequences are well suited for a streamlined and automated conversion to marker-specific Webin checklists. The identification and characterization of plant specimens via the sequencing of specific genome regions (“plant DNA barcodes”; [22]) have become a key method in botanical research [23]. Thousands of DNA sequences have been generated in investigations on suitable plant DNA barcoding markers [24–26], and plant DNA barcoding is now routinely applied across evolutionary, ecological and conservation research [22,23,27], even in regional studies [28–30]. Future investigations applying plant DNA barcoding will invariably require the submission of novel DNA sequences to public sequence repositories [7,27], and a user-friendly, streamlined data submission process can be instrumental to their data sharing process [3]. In fact, plant DNA barcoding sequences lend themselves for the application of software tools that streamline and automate their submission process, because common barcoding markers display (a) a general homogeneity in sequence length and gene synteny, at least within most target lineages [31,32], (b) a general absence of structural inversions or strong secondary structure [32,33], and (c) the presence of conserved regions, preferentially in the flanking parts of the marker, that allow the anchoring of universal PCR primers and, thus, bidirectional sequencing [31–33]. Unfortunately, the preparation process for submissions of DNA sequences that are commonly generated in plant DNA barcoding experiments is insufficiently automated, especially to the sequence database of ENA. A software tool is needed that streamlines and automates the preparation of such barcoding sequences, primarily for submissions via the interactive Webin submission system of ENA.

The plant sciences community would benefit from a software tool that streamlines the generation of marker-specific Webin checklists, which is tedious to conduct by hand and difficult to code dynamically due to the idiosyncrasies of the different checklists (Fig. 1). Specifically, it would be desirable to automate the conversion of EMBL- or GenBank-formatted flat files, which can be generated via various software suites for DNA sequence assembly and annotation (e.g., Geneious, Artemis), into correctly formatted Webin checklists that require nothing more but their upload to ENA via the interactive Webin submission system (Table 1). The software should hereby be available as an open-source tool that can be customized and expanded by other researchers. In the present investigation, we present such a tool. We report about the development and application of a Python package, entitled “EMBL2checklists”, that takes annotated DNA sequences of common plant DNA barcoding regions and associated metadata as input and returns properly-formatted checklists that are ready for data upload to ENA via the interactive Webin submission system (Fig. 2).

**Table 1.** Overview of the bioinformatic steps involved in submitting novel DNA sequence data to ENA while using EMBL2checklists, starting from assembled DNA sequences.

|  | Bioinformatic step | Details of step | Manual processing | Available software |
| --- | --- | --- | --- | --- |
| 1 | Sequence annotation | Sequence features and feature qualifiers using INSDC-compatible keywords are added to each DNA sequence. | high | Geneious<sup>1</sup>, Artemis<sup>1,2</sup> |
| 2 | Saving as flat file | Multiple annotated DNA sequences are saved as a single multi-sequence EMBL- or GenBank-formatted flat file. | minimal | Geneious<sup>1</sup>, Artemis<sup>1,2</sup> |
| 3 | Flat file validation | flat file format, feature table syntax or taxonomic status of organism, among other aspects, are validated. | minimal | EMBL flat file validator<sup>1</sup> (cit.), SBOL validator<sup>2</sup> (cit.) |
| 4 | Conversion to checklist | The EMBL- or GenBank-formatted flat file is converted into the Webin checklist file format. | minimal | EMBL2checklists<sup>1,2</sup> |
| 5 | Upload to ENA | The checklist is uploaded to ENA via the interactive route of the Webin submission system. | minimal | Any web browser |

Table notes. <sup>1</sup>For EMBL-formatted flat files; <sup>2</sup>For GenBank-formatted flat files

**Fig 2.**
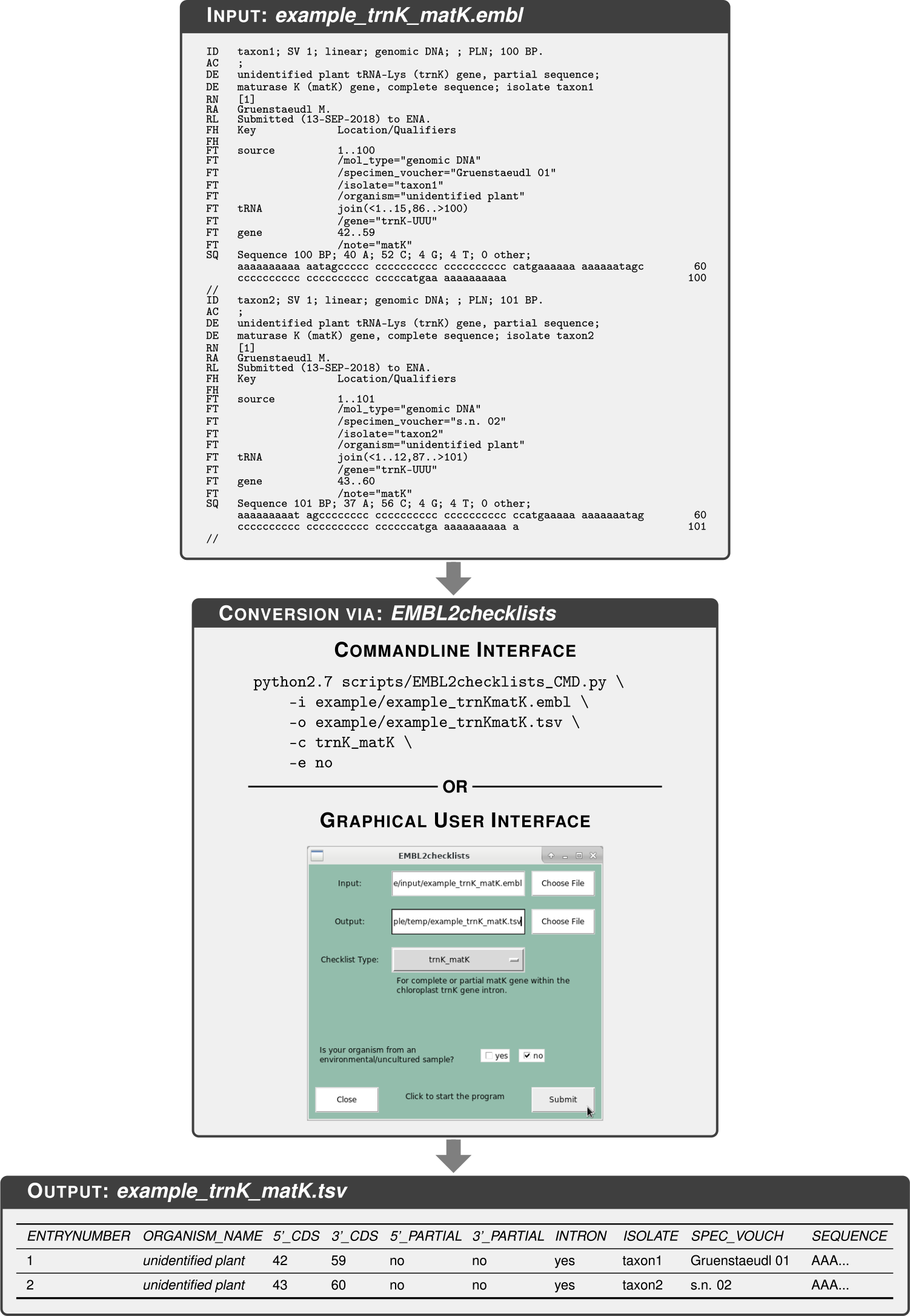
Overview of the conversion process from an EMBL-formatted flat file to a submission-ready Webin checklist via the application of EMBL2checklists. Name and content of the input and output files displayed are identical to the corresponding example test files co-supplied with the software.

## Materials and methods

### Implemented checklist types

The software EMBL2checklists was designed to convert annotated DNA sequences and associated metadata into six different Webin checklist types. Sequence submission via the Webin submission system is primarily conducted via marker-specific checklists [15]. These checklists are pre-tailored to the idiosyncrasies of different genome regions, with different checklists displaying marker-specific customizations in order to capture the distinct information of the genomic regions under study (Fig. 1). A conversion of annotated DNA sequences into Webin checklists, thus, needs to take these marker-specific customizations into account. For example, a checklist that contains sequence information on the plastid *trnK/matK* region will be more complex than a checklist on the nuclear ribosomal external transcribed spacer (ETS) due to the location of the gene *matK* inside the group II intron of the tRNA gene for Lysine (*trnK*-UUU; [34]). Thus, the Webin checklist for *trnK/matK* comprises eight mandatory columns, whereas the Webin checklist for the ETS comprises only five (Table 2). To enable submissions of a wide range of genomic regions to ENA, numerous marker-specific checklists have been implemented in Webin (https://www.ebi.ac.uk/ena/submit/annotation-checklists), but only a small number of these are relevant to plant DNA barcoding. The software EMBL2checklists is designed to convert annotated DNA sequences and associated metadata into one of six different Webin checklists. Given an EMBL- or GenBank-formatted flat file with the correct annotation features and feature qualifiers as input (Table 2), EMBL2checklists can generate marker-specific checklists for a series of DNA markers that are commonly employed in plant DNA barcoding. These markers are: (i) a common gene intron (e.g., *trnL* intron; [35]); (ii) a common intergenic spacer (IGS; e.g., *trnH-psbA*; [36]); (iii) the plastid *trnK/matK* region [37]; (iv) the nuclear ribosomal rRNA-encoding rDNA genes (e.g., 18S rDNA; [38]); (v) the nuclear ribosomal internal transcribed spacer (ITS; [24]); and (vi) the nuclear ribosomal ETS ([39]).

**Table 2.**
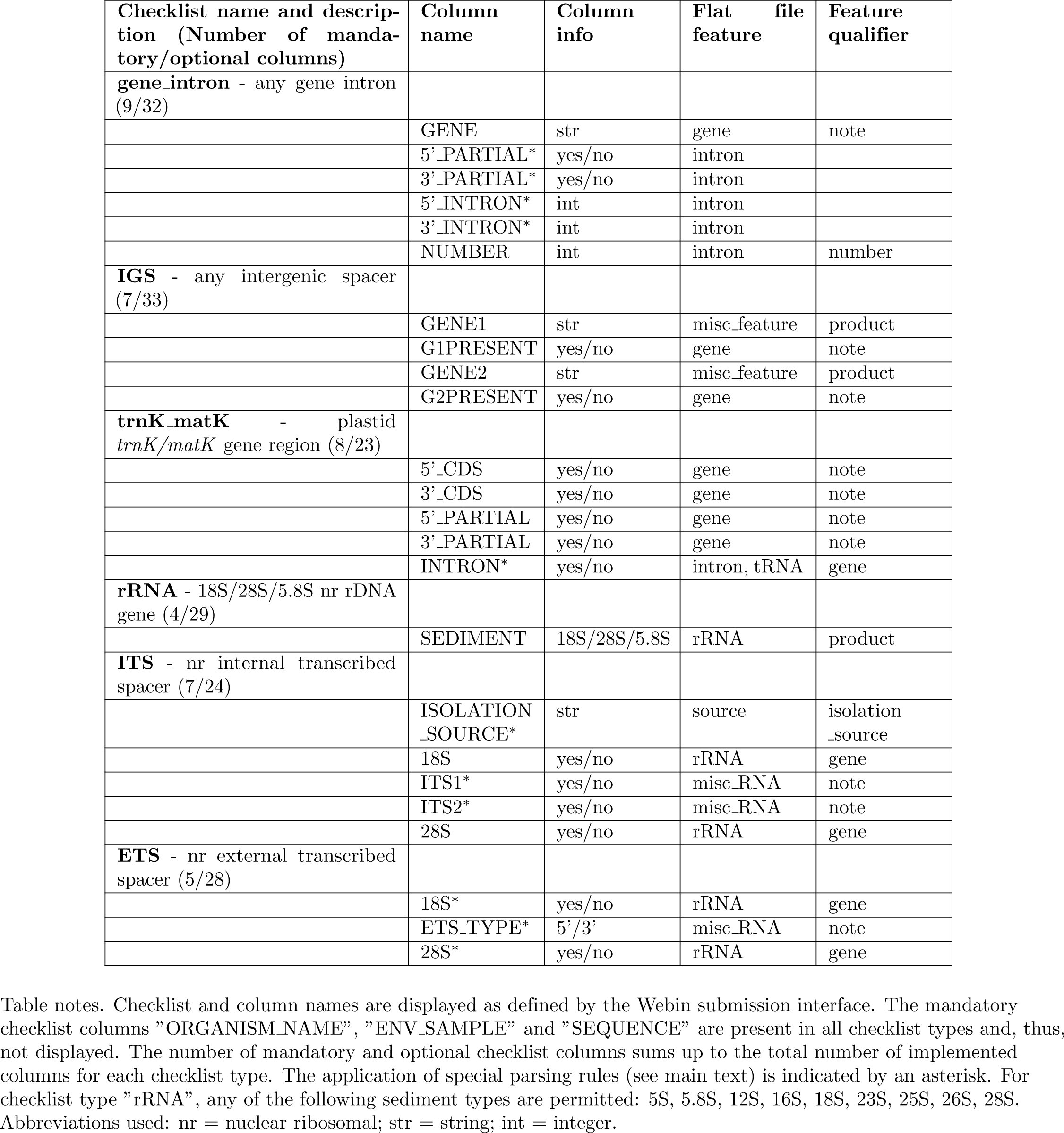
Overview of the mandatory column name, annotation feature and feature qualifier specifications of the six checklist types implemented in EMBL2checklists.

### Conversion of multiple sequence records

EMBL2checklists is able to convert multiple sequence records contained in input flat file into a single Webin checklist. EMBL- or GenBank-formatted flat files may contain multiple sequence records, each with a specific set of annotation features and sequence metadata. EMBL2checklists accepts such a user-selected flat file as input, parses each sequence record individually, and writes the parsed information to the output file. Specifically, the software converts the sequence information contained in each sequence record into a separate line of the resulting checklist. Programmatically, EMBL2checklists parses the flat file via the BioPython library [40] and then iterates through the sequence records, processing one record at a time (Fig. 3). During each iteration, the DNA sequence of a record, its annotation features, and its associated metadata are extracted. The extracted information is formatted to a pre-tailored, tab-delimited spreadsheet, the precise type of which had been selected by the user during program initiation. Eventually, EMBL2checklists collects the information of all successfully processed records and appends it to the output file so that the number of lines written to the checklist equals the number of sequence records successfully parsed from the input. This record-by-record processing of the input file allows the parsing algorithm to evaluate the sequence records individually and to skip specific records in the event of an error.

**Fig 3.**
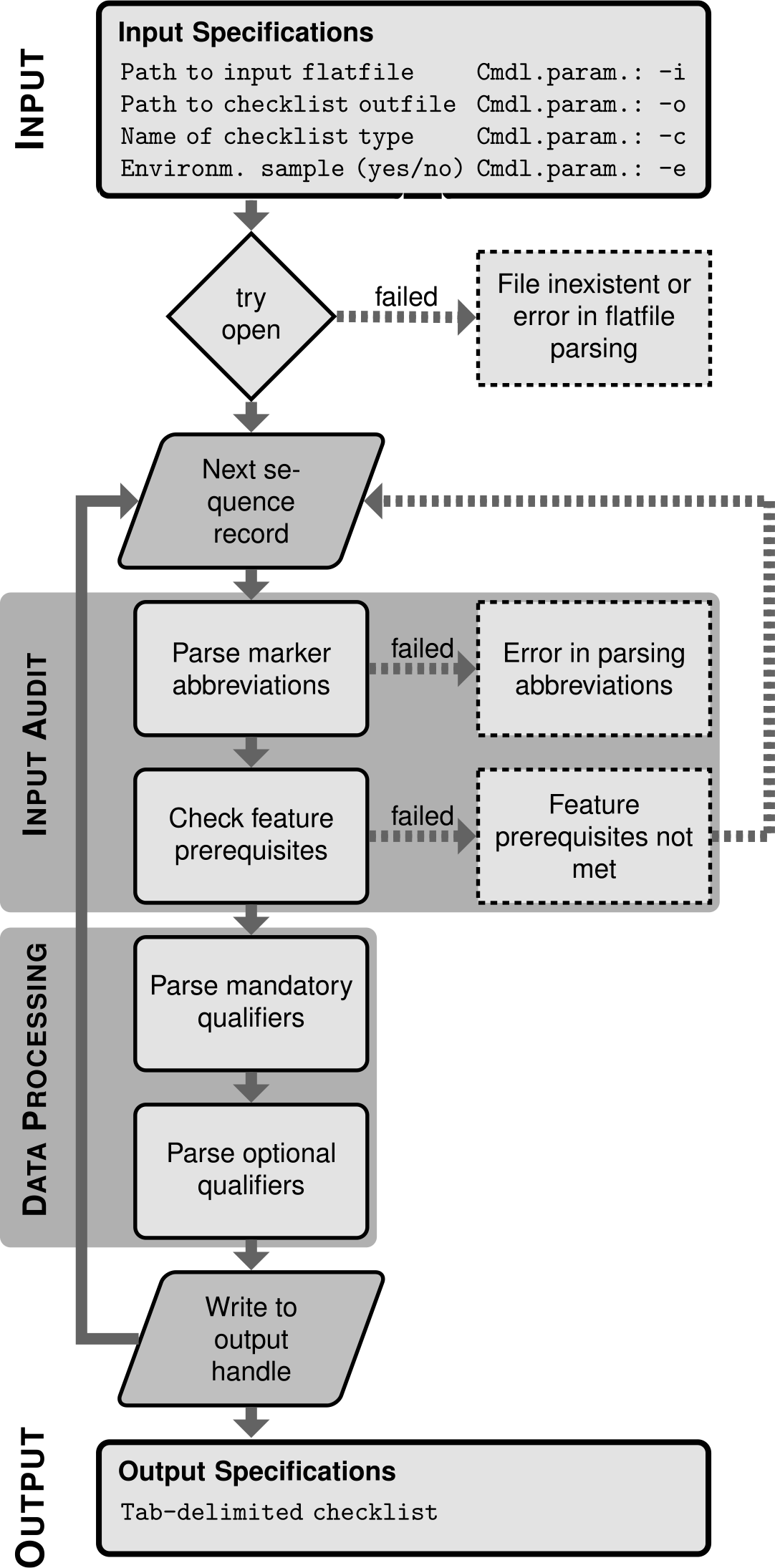
Overview of the internal structure of EMBL2checklists. The overview illustrates the two main processes that are executed sequentially for each sequence record: input audit and data processing. Rhomboid fields indicate the start and end points of the loop across sequence records. Field with dotted outlines indicate states when standard data processing fails.

### Internal structure of software

EMBL2checklists conducts a series of data parsing steps that are executed sequentially for each sequence record and that include evaluations of feature prerequisites and the coherence between input data checklist selection. The structure of EMBL2checklists consists of two main processes that are executed sequentially for each sequence record: input audit and data processing (Fig. 3). Upon initialization, EMBL2checklists evaluates the presence, integrity, and syntax of the input flat file via the data parser of BioPython. Then, it proceeds to input audit and data processing. During input audit, two different checks are conducted on a given sequence record. These checks are implemented as separate functions and comprise the evaluation if (a) the DNA marker abbreviations found among the annotation features and their qualifiers are coherent with the user-selected checklist type (“parse marker abbreviations” in Fig. 3), and (b) the sequence record contains the minimally necessary annotation features to generate a functional checklist (“feature prerequisites”). Thus, the input audit evaluates if a given sequence record contains the necessary annotation features, feature qualifiers and qualifier values for the chosen checklist type. For example, a sequence record of a ribosomal DNA region that does not contain at least one feature of class “rRNA” in the feature table and in which that feature does not contain at least one qualifier of class “product” is flagged as an error because information on the mandatory checklist qualifier “SEDIMENT” cannot be parsed from such a sequence record (Table 2). Sequence records that fail the evaluation of minimal feature prerequisites are skipped, whereas those that fail the parsing of correct marker abbreviations terminate the entire software execution (Fig. 3) because the latter error is indicative of an incorrect checklist selection by the user. Upon successful input audit, data processing of the sequence record is started. During data processing, the annotation features and feature qualifiers of a sequence record relevant to the chosen checklist type are parsed. First, only those qualifiers are parsed that a valid checklist for the selected checklist type could not be generated without (i.e., “mandatory qualifiers”; Fig. 3). Then, qualifiers are parsed, which are permissible but not required for the selected checklist type (i.e., “optional qualifiers”). During each parsing step, the spelling of feature and qualifier names is evaluated against an INSDC-compliant dictionary of feature definitions, which serves two purposes. First, it ensures that feature and qualifier names are spelled and formatted in compliance with the definitions of the INSDC [14]. Second, it ensures the correct transfer of information from the annotation features into the columns of the resulting Webin checklists. If an error is encountered during data processing, the processing of that sequence record fails and raises an exception while communicating a short error message to the user. Upon data processing, the information of the current sequence record is saved into an output handle and the data audit for the next record initiated. Upon processing all sequence records, the writer function of EMBL2checklists appends the parsed information of each sequence record as independent lines to the output file.

### Special parsing rules

EMBL2checklists applies a series of special parsing rules in order to accommodate the idiosyncratic structure and information content of the different checklist types. Data processing is not homogeneous across all checklist types but includes the application of special parsing rules for select checklists. For the checklist type “gene_intron”, for example, the start and end position (“5’JNTRON” and “3’_INTRON” in Table 2) and the completeness of the intron (”5’_PARTIAL” and “3’_PARTIAL”) is determined by the intron location information, not by its qualifier values. For checklist type “trnK_matK”, either an intron feature or a tRNA feature for gene *trnK*-UUU must be present in the sequence record. For checklist type “ITS”, two special rules apply: (a) if environmental samples are processed, the otherwise optional checklist column “ISOLATION_SOURCE” becomes mandatory; and (b) the completeness of the rDNA gene 5.8S is inferred based on the presence of ITS1 and ITS2. For checklist type “ETS”, either a rRNA feature for the 18S and the 28S rDNA gene, or a misc_RNA feature with the info “5’ ETS” or “3’ ETS” must be present in the sequence record. To maintain an accurate implementation of these parsing rules across different development stages, customary checks via the Python “unittest” framework [41] were added to the software.

### Input parameters and output specifications

Upon initializing the software, users of EMBL2checklists must specify four input parameters (Fig. 2): (a) the name of the input flat file; (b) the name of the checklist output file; (c) the type of checklist selected by the user, and (d) a statement if the sequence records classify as environmental samples. Input parameter (a) must contain the name of, and the file path to, an EMBL- or GenBank-formatted flat file that comprises one or more sequence records. The precise file format of the input flat file is automatically identified based on the file ending (i.e., “.embl” or “.gb”). The structure of each sequence record must be compliant with the identified flat file format and consist of a multi-line feature table, followed by the interleaved DNA sequence. The source feature, which pertains to the sequence as a whole and often contains sequence metadata, must be located at the top of the feature table. Annotation features, which indicate the type and boundaries of localized sequence features, must be located below the source feature. Input parameter (b) must contain the name of, and the file path to, the output checklist. The checklists generated by EMBL2checklists are human-readable and can be edited by any common spreadsheet editor such as Microsoft Excel (Microsoft Corporation, Redmond, WA, USA) or LibreOffice (The Document Foundation, Berlin, Germany). If all sequence records of the input file are processed correctly, the output checklist displays as many lines as the number of sequence records in the input file (disregarding the title line of the checklist). Input parameter (c) must be the name of one of six checklist types implemented in EMBL2checklists and specifies the rule set applied during input audit and data parsing of the sequence records. Input parameter (d) must be either “yes” or “no” and specifies the classification of all sequence records of an input file as environmental samples. This parameter is implemented in EMBL2checklists because its information is mandatory for certain Webin checklist types. It is typically answered with “yes” if the DNA sequences under study were generated as part of a metabarcoding experiment. Upon specifying each of these parameters, EMBL2checklists begins to process the input file.

### Commandline and graphical user interface

EMBL2checklists was developed for classical plant biologists and bioinformaticians alike. Thus, the software is equipped with a graphical user interface (GUI) as well as a command-line interface (CLI) for its operation (Fig. 2). The GUI is based on the Python library “Tkinter” [42] and designed to provide an intuitive and easy-to-use interface that allows users with little or no bioinformatics knowledge to operate the software. To execute EMBL2checklists via the GUI, users enter the four input parameters via the available input fields and drop-down menus. To receive help and detailed explanations via the GUI, users may hover their mouse pointer over a field of interest, which initiates a help bubble next to the pointer. Moreover, details of individual checklist types are automatically displayed upon selecting one of the implemented checklists from the drop-down menu. If EMBL2checklists raises an exception while operating under the GUI, an error message is printed to a pop-up window of the GUI. The GUI can be accessed through file “EMBL2checklists_GUI.py” of the scripts folder. More information on the design and functionality of the GUI of EMBL2checklists is available in [43]. The CLI employs functions of the Python library “argparse” [44] and allows more experienced users to execute the software via the command-line and to integrate the software into larger bioinformatic workflows. To execute EMBL2checklists via the GUI, users specify the name of the input flat file via command-line argument “-i”, the name of the checklist output file via argument “-o”, the type of checklist via argument “-c”, and the classification of the sequence records as environmental samples via argument “-e”. To receive help and detailed explanations, CLI users may invoke command-line argument “-h” (i.e., “python2 script/EMBL2checklists_CLI.py-h”). If EMBL2checklists raises an exception while operating under the CLI, an error message is printed to the standard output stream. The CLI can be accessed through file “EMBL2checklists_CLI.py” of the scripts folder.

### Release, installation and operation

EMBL2checklists was written in Python 2.7 [45] and is, thus, platform independent. It can be executed on any system equipped with a Python 2 compiler and after the installation of the necessary Python dependencies. The software uses three separate Python packages as dependencies: Biopython [40], argparse [44] and Tkinter [42]. EMBL2checklists is open source and released under the BSD 3-Clause license (https://opensource.org/licenses/BSD-3-Clause). Other bioinformaticians are, thus, able to expand and customize the software to fit their own data submission projects. EMBL2checklists is available via the Python Package Index (https://pypi.org/project/EMBL2checklists/) and can be installed via any PyPI-compatible package management system such as pip (https://pip.pypa.io) or setuptools (https://pypi.org/project/setuptools/). For example, users may type the following command to install EMBL2checklists on their system:

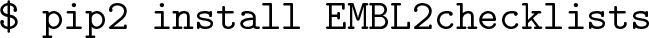

EMBL2checklists was successfully tested on a Windows (Microsoft Windows 7), a MacOS (MacOS 10.12.6 - Sierra) and two different Linux environments (Arch Linux 4.18.9 and Debian 9.0). For a typical execution of EMBL2checklists via the CLI, a user may type the following command:

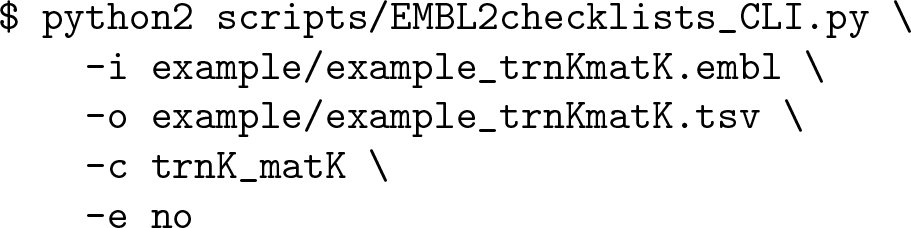

## Results and Discussion

### Application of software on empirical data

The utility of EMBL2checklists to plant biology is illustrated by its application in the submission process of DNA sequences to ENA by three recent investigations. Specifically, EMBL2checklists was employed for the preparation of hundreds of DNA sequences used in two plant phylogenetic and one fungal metagenomic investigation [46–48]. The two plant phylogenetic investigations utilized common plant DNA barcoding markers to infer the phylogenetic history of select plant lineages [46,47]. The fungal metagenomic investigation utilized nrDNA barcodes to characterize arbuscular mycorrhizal soil fungi [48]. In each case, EMBL2checklists was used to convert flat files in GenBank format that were generated from sets of assembled and annotated sequences via the software suite Geneious. Upon conversion to checklists, the sequence data was uploaded to ENA via the interactive Webin submission system, and accession numbers were received from ENA by email within less than 48 hours of submission.

### Post-processing of checklists and work-sharing

Due to the data format of Webin checklists and the structure of the interactive Webin submission process, the use of EMBL2checklists for sequence submissions to ENA displays two special advantages: it allows the simple post-processing of checklists, and it enables work-sharing during the submission process. First, the data format of Webin checklists is human-readable and, thus, allows the manual addition of column information, should users wish to augment the checklist prior to submission. Specifically, the easily-accessible checklist data structure allows users to modify or append column content, as long as the name and order of existing columns remain unchanged. For example, users who wish to add information about the geographic location of one or more sequences after having processed the sequence data with EMBL2checklists can simply add a column entitled “LOCALITY” before column “SEQUENCE” of the checklist and add geographic location information for one or more sequences. Likewise, users who wish to combine multiple checklists can do so, as long as the number and order of columns are identical (and the title line of the second checklist is removed). Second, EMBL2checklists allows the implementation of a work-sharing strategy during the preparation of sequence submissions because the information contained within the identification, description and reference lines of a sequence record (such as author name or author institution) is not saved as part of the checklist output. The software only processes the feature table and the DNA sequence of a sequence record. Ancillary information of a dataset must be associated with the sequence data during the interactive submission process. Specifically, personal and institutional information of the submitter is associated with the data through the Webin submission service prior to data upload, irrespective of the checklist type. Hence, EMBL2checklists does not need to be executed by the same person that conducts the data upload or has generated the sequence but allows a work-sharing strategy in which one person (or section of a workflow) conducts the data conversion via EMBL2checklists, while another person (or section of a workflow) conducts the data submission. Work-sharing may be helpful if the sequence submission process is centralized within a lab or academic institution, allowing those researchers that prepare the data for submission to ENA to be different from those that actually conduct the data upload.

### Data converters and other ENA submission strategies

The paucity of file formats acceptable for data submission to public sequence databases is one of the main bottlenecks in the effort to increase participation in public data sharing and has spurred the recent development of various data converters. The software EMBL2checklists is one of several current projects that aim to provide automated data conversion between the EMBL or GenBank flat file format and data formats that are commonly parsed by biological software and databases [20, 49, 50]. The underlying aim of many of these projects is to simplify the conversion process of sequence data into file formats that are accepted during submissions to public sequence databases [20, 49]. Given the custom validation criteria and the idiosyncratic submission procedures employed by many of these databases, such data converters represent an important means to enable user-friendly data submissions, at least until such time as submission procedures across INSDC databases are standardized [51]. EMBL2checklists was specifically designed to bridge the gap between common software suites for DNA sequence assembly and annotation (e.g., Artemis, Geneious) and the interactive data submission process of ENA. Compared to most other recent data converters, EMBL2checklists supports the conversion of flat files that contain multiple sequence records. At the same time, EMBL2checklists aims to fulfill a seemingly counterintuitive task: The software converts EMBL- or GenBank-formatted flat files into Webin checklists, which the receiving database of ENA eventually converts back into the flat file format. Theoretically, it would be easier to upload annotated DNA sequences in EMBL flat file format to ENA directly, but presently only the Entry Upload service (https://ena-docs.readthedocs.io/en/latest/prog_12.html) and, to lesser extent, the Webin command-line submission interface (https://github.com/enasequence/webin-cli) support flat file submissions of this data type. In practice, both of these programmatic services exclude a considerable number of regular users aiming to upload annotated DNA sequences to ENA due to the bioinformatics expertise required. Until methods are in place that allow a more user-friendly upload of annotated DNA sequences in flat file format to ENA, a back-conversion to checklists as conducted by our software remains the most practical solution. Thus, EMBL2checklists currently represents the most user-friendly option for plant biologists without bioinformatics experience to automatically generate submission-ready checklists from common plant DNA sequence data.

## Conclusion

The lack in automated conversion between EMBL- or GenBank formatted flat files and submission-ready Webin checklists represented a gap that compelled many researchers to conduct manual data processing before submitting data to the public sequence database ENA. By developing the software EMBL2checklists, we have filled this gap. EMBL2checklists is designed as an easy-to-use software application that bridges the gap between common software suites for DNA sequence assembly and annotation and the interactive data submission process of ENA. The software converts annotated DNA sequences plus associated metadata into properly-formatted Webin checklists for submission to ENA. Specifically, the software takes the idiosyncrasies of marker-specific checklist types into account and generates submission-ready checklists for specifically those DNA markers that are commonly employed in plant DNA barcoding. Thus, EMBL2checklists can be employed to prepare the most common plant DNA barcoding marker sequences for upload and submission to ENA via the interactive Webin submission system. Users may generate input files for our software through any of several common sequence analysis environments such as Artemis or Geneious and then employ the GUI of EMBL2checklists to prepare their sequence submissions. Upon processing with EMBL2checklists, the user receives a checklist that can be directly uploaded to ENA or further edited with a common spreadsheet editor. The utility of EMBL2checklists is best illustrated by its application during the submission preparation of hundreds of DNA sequences submitted to ENA during the publication process of several recent investigations [46–48]. With the development of EMBL2checklists, we hope to provide a useful software tool to plant biologists and bioinformaticians alike, increase the amount of sequence data deposited to public sequence databases [7] and advance the idea of publicly-shared research data [9,10]. By extension, we believe that EMBL2checklists may play an important role in future data management and data stewardship of plant DNA sequence data under the FAIR data principle [52,53].

## Acknowledgments

The authors would like to acknowledge the high-performance computing service of the ZEDAT of the Freie Universitat Berlin for providing allocations of computing time. The development of a GUI for the presented software constitutes part of a thesis by YH toward a bachelor of science degree.

## Data availability statement

EMBL2checklists is available free of charge via the Python Package Index (https://pypi.org/) under https://pypi.org/project/EMBL2checklists/ and can be installed via any PyPI-compatible package management system such as pip (https://pip.pypa.io) or setuptools (https://pypi.org/project/setuptools/). The source code of EMBL2checklists is available via the GitHub page of MG under https://github.com/michaelgruenstaeudl/EMBL2checklists.

## Author contributions

MG conceived the software, wrote its core functions and generated an alpha version of the software; YH added additional functions to the software and wrote its GUI; MG applied the software on the empirical data; MG wrote the manuscript.

